# Suppression of p53 response by targeting p53-Mediator binding with a stapled peptide

**DOI:** 10.1101/757401

**Authors:** BL Allen, K Quach, CB Levandowski, JD Rubin, T Read, RD Dowell, A Schepartz, DJ Taatjes

## Abstract

DNA-binding transcription factors (TFs) remain challenging to target with molecular probes. Many TFs function in part through interaction with Mediator; we sought to block p53 function by disrupting the p53-Mediator interaction. Through rational design and activity-based screening, we characterized a stapled peptide, with functional mimics of both p53 activation domains, that selectively inhibited p53- and Mediator-dependent transcription *in vitro*. This “bivalent peptide” also suppressed p53 transcriptional response in human cancer cells. Our strategy circumvents the TF and instead targets the TF-Mediator interface, with desired transcriptional outcomes. Different TFs target Mediator through different subunits, suggesting this strategy could be broadly applied.

## Introduction

Sequence-specific, DNA-binding transcription factors (TFs) drive myriad physiological processes and their mutation or disruption underlies many human diseases (Lee and Young 2013). They are unquestionably high-impact targets for molecular therapeutics. Unfortunately, TFs have proven difficult to target with small molecules (Bradner et al. 2017); their DNA-binding domains are charged and similar to other TFs, and their activation domains are typically unstructured and intrinsically disordered.

Among the estimated ∼1600 TFs in the human genome (Lambert et al. 2018), p53 stands out for its general importance in cancer biology (Khoo et al. 2014; Kruiswijk et al. 2015; Kastenhuber and Lowe 2017). Across many cell lineages, p53 functions as a tumor suppressor and can paradoxically function as an oncogene if it acquires specific “gain-of-function” mutations (Freed-Pastor and Prives 2012); p53 also plays key roles in mammalian development, aging, and stem cell biology. Like many TFs, the p53 protein possesses a DNA-binding domain and an activation domain (AD). The p53AD actually consists of two separate but closely-spaced domains, called AD1 (residues 14 – 26) and AD2 (residues 41 – 57). Whereas most transcriptional activation function can be attributed to p53AD1 (Jimenez et al. 2000; Johnson et al. 2005), loss-of-function p53AD1 mutations retain some ability to activate specific subsets of p53 target genes, and mutation of both AD1 and AD2 is required to mimic a p53-null phenotype (Brady et al. 2011; Jiang et al. 2011).

The human Mediator complex contains 26 subunits and is generally required for RNA polymerase II (pol II) transcription (Poss et al. 2013). Mediator interacts extensively with the pol II enzyme and regulates its function in ways that remain poorly understood; however, a basic aspect of Mediator function is to enable TF-dependent activation of transcription (Poss et al. 2013). In fact, Mediator was originally discovered in *S. cerevisiae* using an *in vitro* assay to screen for factors required for TF-dependent transcription (Flanagan et al. 1991); similar functions have been confirmed for mammalian Mediator complexes (Fondell et al. 1996). Because TFs do not bind pol II directly, it appears that they communicate their pol II regulatory functions indirectly, through the Mediator complex. Upon binding Mediator, TFs induce conformational changes that remodel Mediator-pol II interactions to activate transcription (Taatjes et al. 2002; Meyer et al. 2010). The p53 TF binds Mediator and this interaction has been shown to activate p53 target gene expression *in vitro* and in cells (Ito et al. 1999; Meyer et al. 2010). Oncogenic mutations in p53AD1 disrupt p53-Mediator interactions (Ito et al. 1999) and this correlates with loss of p53 function (Lin et al. 1994). Whereas specific residues and structural details remain unclear, the p53-Mediator interface appears to involve the MED17 subunit (Ito et al. 1999; Meyer et al. 2010). Interestingly, other TFs (e.g. SREBP or nuclear receptors) activate transcription through interactions with different Mediator subunits (Poss et al. 2013).

As selectively targeting TF activation domains has not proven to be a viable strategy to control TF function, here we sought to test whether the same outcome could be achieved by targeting Mediator instead. We chose p53 as a test case because it is well-studied, biomedically important, and contains a well-characterized activation domain. An initial obstacle was that Mediator is large (1.4 MDa, 26 subunits) and its p53 interaction site is not precisely defined. However, we reasoned that the p53 activation domain (residues 13 – 60) evolved to selectively interact with Mediator with high affinity; consequently, we used the native p53AD structure and sequence as a starting point, rather than screen thousands of drug-like compounds. To directly assess Mediator targeting, we used a defined *in vitro* transcription system that recapitulated p53- and Mediator-dependent transcription.

## Results and Discussion

### An in vitro assay to test p53-activated vs. basal transcription

To screen peptides for the ability to selectively block p53-dependent transcription, we required an assay that enabled p53-dependent activation but that could also support basal (i.e. activator-independent) transcription. We previously established an *in vitro* transcription assay using highly purified human factors (Knuesel et al. 2009). A key feature of this assay was that both activated and basal transcription could be reconstituted on naked DNA templates (i.e. DNA templates not assembled into chromatin). To adapt this assay for purposes of measuring basal vs. p53-activated transcription, we generated templates with Gal4 DNA binding sites upstream of the adenovirus major late promoter sequence (Fig. 1A). Upon titration of a Gal4 DNA Binding Domain (DBD)-p53 Activation Domain (AD; residues 1-70) fusion protein into this system, we observed pol II-dependent transcription that was dependent on p53AD and Mediator (Fig. 1B **and** C). Reactions containing Gal4-p53AD generally produced about two- to four-fold more transcripts compared to reactions with no activator (Fig. 1C). Because experiments without Gal4-p53AD produced a low level of “basal” transcription that could be quantitated, this system allowed assessment of both p53-activated and basal transcription.

**Figure 1.**
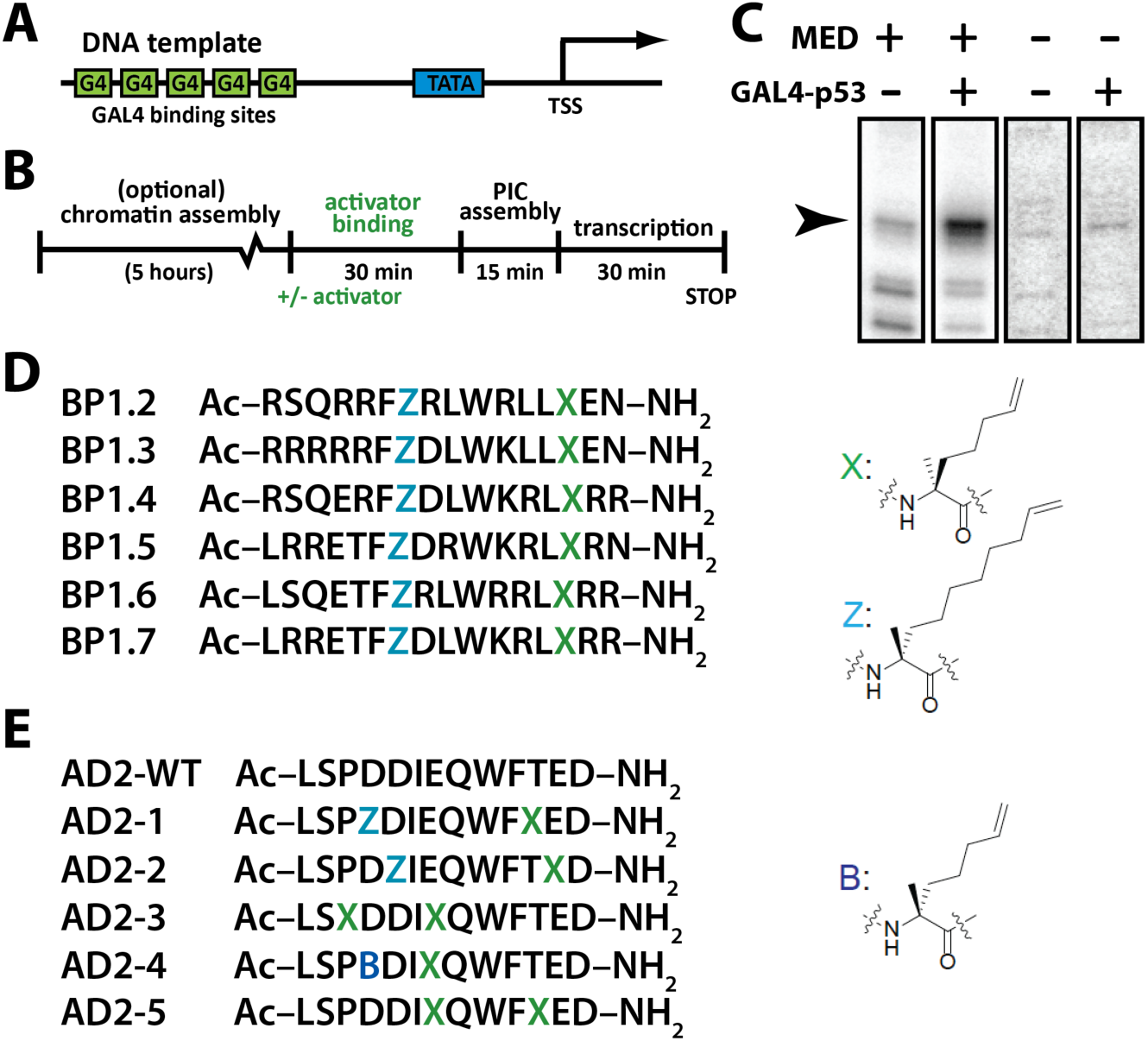
*In vitro* screening protocol and stapled p53AD1 or p53AD2 peptides. (*A*) Promoter DNA template scheme. (*B*) Overview of reconstituted *in vitro* transcription assay (chromatin assembly optional; PIC = Pre-Initiation Complex: TFIIA, TFIIB, TFIID, TFIIE, TFIIF, TFIIH, Mediator, and pol II). (*C*) Representative *in vitro* transcription data from naked DNA templates, showing p53- and Mediator-dependence. (*D*) Sequences of p53AD1 peptides containing diverse penta-arg motifs. Residues Z and X represent α,α-disubstituted amino acids with olefin tethers for hydrocarbon-stapling. (*E*) Sequences of p53AD2 peptides. Residue B represents an α,α-disubstituted amino acid with olefin tethers for hydrocarbon-stapling.

### Design and synthesis of stapled peptides

We designed hydrocarbon-stapled peptide mimetics of the AD1 and AD2 regions of p53. Hydrocarbon-staples were employed to promote helicity within our peptides. Hydrocarbon-stapled peptides have previously been developed to mimic the α-helical portion of p53AD1 that binds MDM2/MDM4 with the goal of blocking the p53-MDM2/MDM4 interaction and restoring wild-type p53 activity (Bernal et al. 2007; Brown et al. 2013). Indeed, ALRN6924, a hydrocarbon-stapled p53AD1 peptide mimetic developed from dual MDM2/MDM4 inhibitor ATSP-7041 (Chang et al. 2013), is being evaluated in clinical trials (ClinicalTrials.gov identifier: NCT02264613 and NCT02909972). For the p53AD1 mimetics, we synthesized N-acetylated versions of the penta-arg-containing peptides BP1.2 - BP1.7 (Quach et al. 2018), which are based on residues 14-29 of p53 (Fig. 1D). This panel of peptides contains an *i, i*+7 hydrocarbon staple at positions 20 and 27 and five arginine residues grafted into various positions, which were originally introduced to improve the cytosolic access of the peptides. For the p53AD2 mimetics, we designed a panel of hydrocarbon-stapled peptides that varied the length and position of the hydrocarbon staple and spanned residues 45-57 of p53 (Fig. 1E). The panel included two peptides with an *i, i*+7 hydrocarbon staple (AD2-1 and AD2-2), two peptides with an *i, i*+4 staple (AD2-3 and AD2-4), and one peptide with an *i, i*+3 staple (AD2-4). Furthermore, both stapled and unstapled variants of the p53AD2 peptides were generated.

### Functional screening of stapled peptide mimics of p53AD1 and p53AD2

Starting with the stapled p53AD1 peptides, we tested whether any would block p53-dependent transcription activation without inhibiting basal transcription. Initial screens were completed with 5 *µ*M of each peptide (BP1.2 – BP1.7; Fig. 1D). At this concentration, all peptides significantly reduced p53 activated transcription, but only BP1.4 and BP1.5 did not also negatively affect basal transcription (Supplemental Fig. 1A). In follow-up experiments, we observed that the BP1.5 peptide was less selective (i.e. negatively affected both p53-activated and basal transcription) compared with the BP1.4 peptide (Supplemental Fig. 1B). We therefore chose the BP1.4 peptide for further testing.

To determine a concentration range in which the BP1.4 peptide selectively blocked p53-activated transcription and not basal transcription, we titrated BP1.4 into transcription reactions at concentrations between 0.9 *µ*M and 9 *µ*M (Supplemental Fig. 1C). Interestingly, BP1.4 activated basal transcription at concentrations of 4 *µ*M and above, which could reflect weak binding of BP1.4 to Mediator (i.e. mimicking p53AD) to promote transcription activation. Although basal transcription was inhibited at the 9 *µ*M titration point, the weak BP1.4-dependent activation made the determination of the IC_50_ for basal transcription impossible using an inhibitor response curve. The IC_50_ describing the inhibition of p53-activated transcription by BP1.4 was 3.2 ± 0.2 *µ*M (Supplemental Fig. 1D). The concentration window in which BP1.4 selectively blocked activated transcription was therefore relatively narrow.

We next tested the p53AD2 peptides (Fig. 1E) in a similar manner. In contrast to the p53AD1 peptides, the p53AD2 peptides either had no effect on p53-activated transcription or non-specifically inhibited both p53-activated and basal transcription at 5 *µ*M (data not shown). Testing further at different peptide concentrations (i.e. increasing concentration if no activity was observed at 5 *µ*M or decreasing concentration if both activated and basal transcription were inhibited) did not reveal any stapled p53AD2 peptides with specificity for p53-activated transcription. These results were not entirely unexpected, as p53AD2 appears to play a lesser role (vs. p53AD1) in activation of p53 target genes *in vivo* (Jimenez et al. 2000; Johnson et al. 2005; Brady et al. 2011; Jiang et al. 2011).

### A bivalent peptide selectively blocks p53-dependent activation in vitro

We hypothesized that covalently linking two peptides with low-to-moderate affinity and specificity could generate a cooperatively binding “bivalent” peptide with improved ability to inhibit p53-dependent activation. Given the inactivity of the p53AD2 peptides tested (Fig. 1E), we elected to tether the BP1.4 peptide to the wild type p53AD2 sequence. In this way, we hoped to generate a competitive inhibitor of p53AD-Mediator binding by recapitulating the combined landscape of p53AD1/AD2 interactions. And because the p53AD1 portion was stapled (i.e. BP1.4), this “bivalent peptide” might effectively compete with WT p53 for Mediator binding.

We synthesized and tested three bivalent peptides (BP1.4 + p53AD2 sequence) that contained a 2-, 6- or 10-unit polyethylene glycol (PEG) linker (bivalent peptide 1, 2, or 3; Supplemental Fig. 2A). Notably, the bivalent peptides were significantly more potent inhibitors of p53-activated transcription than BP1.4 alone. As shown in Supplemental Figure 2 B, bivalent peptides (500 nM) containing either a 6- or 10-unit PEG linker (i.e. bivalent peptide 2 or 3) inhibited p53-activated but not basal transcription. By contrast, the bivalent peptide with a 2-unit PEG linker (i.e. bivalent peptide 1) did not inhibit transcription at 500 nM (Supplemental Fig. 2B). Because the bivalent peptide containing a 6-unit PEG linker (i.e. BP1.4_PEG6_p53AD2; bivalent peptide 2) was easier to synthesize (vs. 10-unit PEG), it was used for all future experiments. For simplicity, this molecule (bivalent peptide 2, Supplemental Fig. 2A) will be called the “bivalent peptide” throughout this paper.

**Figure 2.**
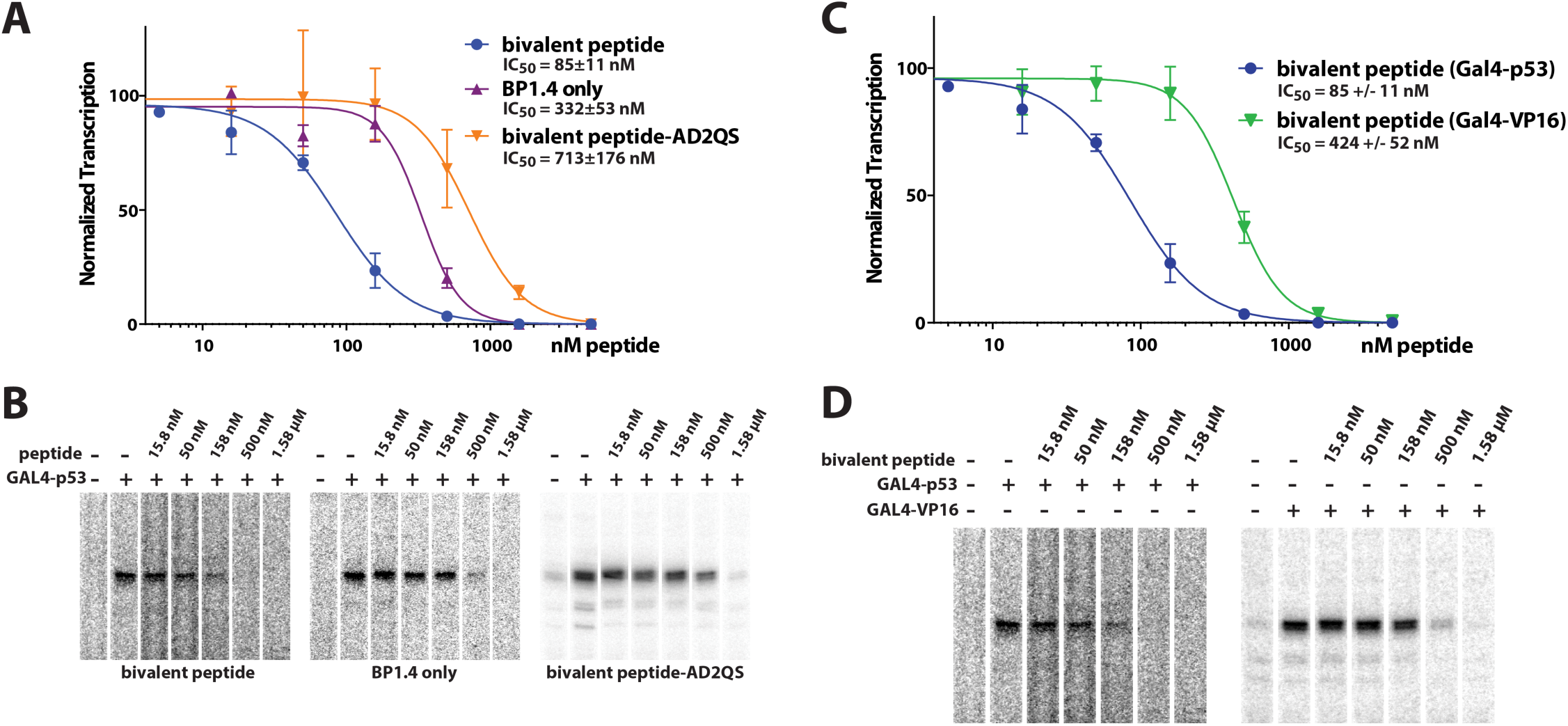
*In vitro* transcription on chromatin templates reveals bivalent peptide is a potent and selective inhibitor of p53-dependent transcription activation. (*A*) IC_50_ plot showing activity of bivalent peptide (n = 3 to 8) vs. BP1.4 (stapled p53AD1 mimic; n = 2 to 6) or a bivalent peptide with a mutated p53AD2 region (n = 3 to 9). The reduced activity of BP1.4 and the AD2 mutant indicate that both p53 activation domains contribute to bivalent peptide function. (*B*) Representative data from experiments plotted in *A*. (*C*) IC_50_ plot showing that bivalent peptide is selective for p53; repressive activity is reduced with GAL4-VP16 (n = 2 to 6), which binds a different Mediator subunit compared with p53. (*D*) Representative data from experiments plotted in *C*. Vertical lines in plots represent standard error of the mean (panel A, C).

The *in vitro* transcription assays on naked DNA templates demonstrated improved potency of the bivalent peptide and also confirmed that it inhibited p53-activated transcription but not basal transcription. We next tested its function on more physiologically relevant chromatin templates, in which basal transcription is repressed. In fact, a TF activation domain (such as p53AD) and Mediator are required for transcription on chromatin templates (Naar et al. 1998; Meyer et al. 2010), due to the ability of Mediator to relay the activation signal from the TF directly to the pol II enzyme (Poss et al. 2013). *In vitro* transcription assays with chromatin templates revealed that the bivalent peptide had a half maximal inhibitory concentration (IC50) of 85 nM when added to reactions activated by Gal4-p53AD. By contrast, the BP1.4 peptide alone had an IC_50_ of 330 nM in these assays, about five-fold worse than the bivalent peptide (Fig. 2A, B). Upon introduction of a QS mutation into p53AD2, which blocks its activation function *in vivo* (Jiang et al. 2011), the IC_50_ increased to 713 nM, about ten-fold higher than the bivalent peptide and two-fold higher than BP1.4 alone (Fig. 2A, B). Collectively, these results suggest that both p53AD1 and p53AD2 contribute to Mediator-dependent transcriptional activation *in vitro*.

Finally, we assessed whether the bivalent peptide would selectively block p53-dependent transcription compared with VP16, a viral activation domain. Whereas p53 and VP16 both interact with Mediator, they do so through different subunits (MED17 and MED25, respectively) (Ito et al. 1999; Milbradt et al. 2011; Vojnic et al. 2011). In contrast to Gal4-p53AD (85 nM), the bivalent peptide had an IC_50_ of 424 nM in the presence of Gal4-VP16 (Fig. 2C, D). These data, which resulted from identical DNA templates assembled into chromatin, suggested that the bivalent peptide selectively blocked the p53–Mediator interaction vs. the VP16–Mediator interaction; thus, p53-dependent transcription activation was more effectively inhibited compared with VP16-dependent transcription.

### Bivalent peptide inhibits p53AD-Mediator interaction in vitro

To further test whether the bivalent peptide would selectively block the p53AD–Mediator interaction, we performed a series of biochemical experiments, as outlined in Supplemental Figure 3 A. The p53AD can bind Mediator with specificity and apparent high affinity (Ito et al. 1999); for example, the p53AD itself (residues 1-70) is sufficient to selectively isolate Mediator from partially purified cell extracts (Meyer et al. 2010). As shown in Supplemental Figure 3 B, p53AD binding to Mediator was markedly reduced (approximately 60% bound vs. no peptide controls) in the presence of the bivalent peptide. These data are consistent with *in vitro* transcription results (Fig. 2) and reveal that the bivalent peptide functions, at least in part, by blocking p53AD–Mediator interactions.

By using purified factors, the *in vitro* biochemical data (Supplemental Fig. 3 & Fig. 2) establish direct inhibition of p53AD–Mediator binding by the bivalent peptide. ChIP assays were considered to further assess inhibition of Mediator recruitment in cells; however, limited available quantities of the bivalent peptide precluded a rigorous analysis. Because the genomic occupancy of Mediator correlates with pol II recruitment and transcription, we instead turned to RNA-Seq, which requires fewer cells and less optimization, to test bivalent peptide function in cells.

**Figure 3.**
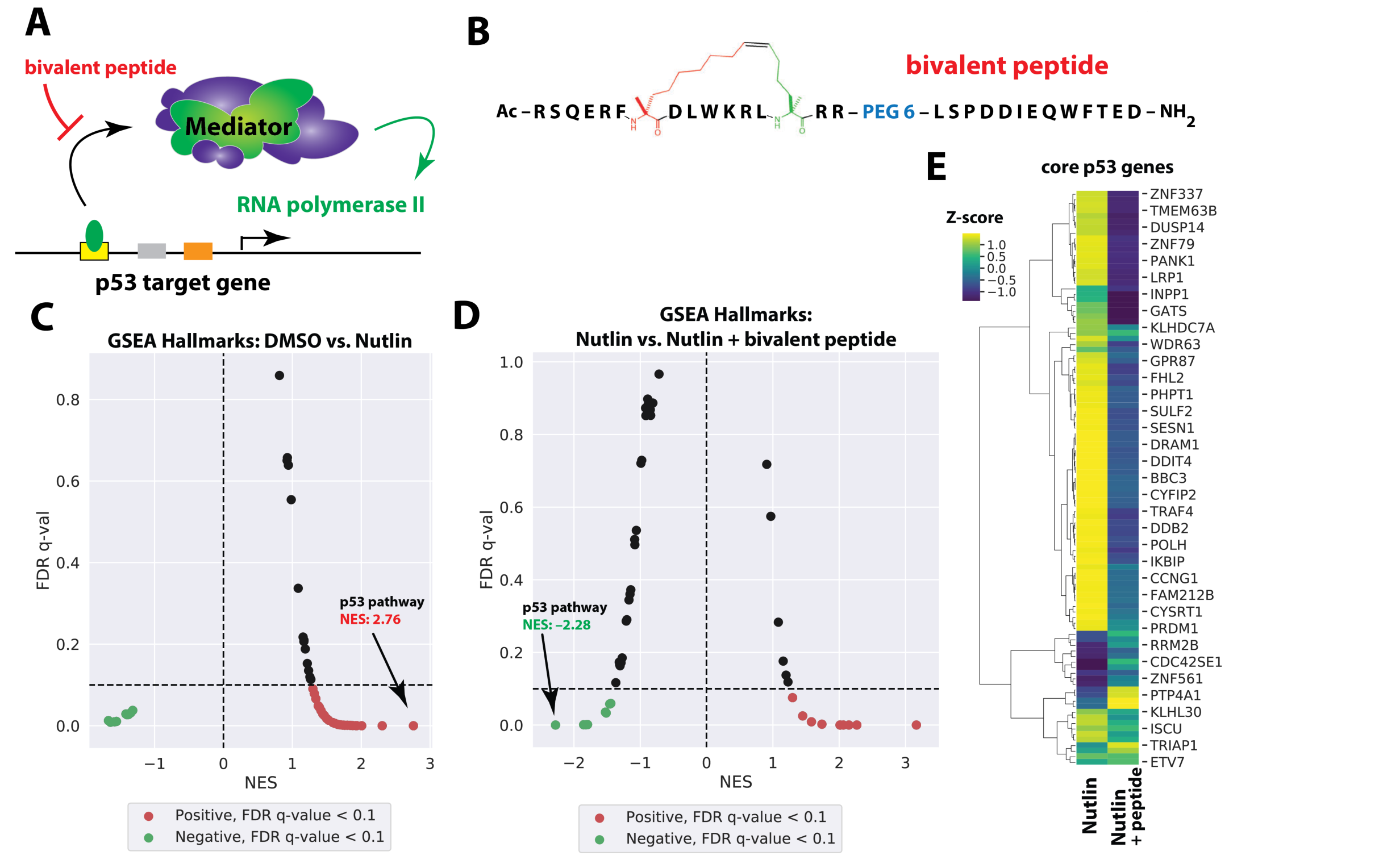
Bivalent peptide blocks p53 response in Nutlin-treated HCT116 cells. (*A*) Simplified schematic showing Mediator recruitment and activation of pol II function via p53. (*B*) Structure of bivalent peptide. (*C*) GSEA hallmarks moustache plot showing robust p53 pathway activation upon treatment with Nutlin-3a (10 *µ*M, 3h). (*D*) GSEA hallmarks moustache plot showing effect of bivalent peptide in Nutlin-treated cells. Note the p53 pathway shows negative enrichment in cells treated with the bivalent peptide. (*E*) Heat map showing relative expression of core set of 103 p53 target genes (Andrysik et al. 2017) in control (DMSO) vs. Nutlin-treated cells, in absence or presence of bivalent peptide. In agreement with GSEA plots from panels C & D, Nutlin-3a induction of p53 target genes is reduced in cells treated with bivalent peptide.

### Bivalent peptide reduces activation of p53 targets in Nutlin-stimulated HCT116 cells

Prior analysis of the BP1.4 peptide showed that it is not effectively taken up by cells (Quach et al. 2018), and given its larger size, the bivalent peptide was expected to have poor cellular uptake. To circumvent this issue, we used a well-tested electroporation protocol to enhance cell uptake of the bivalent peptide (see Methods). HCT116 cells were electroporated either in the presence of bivalent peptide or vehicle (water), with or without Nutlin-3a (Fig. 3A, B). Nutlin-3a is a small molecule that activates and stabilizes p53 by inhibiting MDM2, a repressor of p53 (Vassilev et al. 2004). A 3h treatment time was used based upon experiments that suggested that the bivalent peptide was biologically active for only a limited time in cells (see Methods). After 3h Nutlin-3a treatment (or DMSO control, ± bivalent peptide), nuclear RNA was isolated and biological replicate RNA-Seq libraries were prepared (Supplemental Fig. 4A).

**Figure 4.**
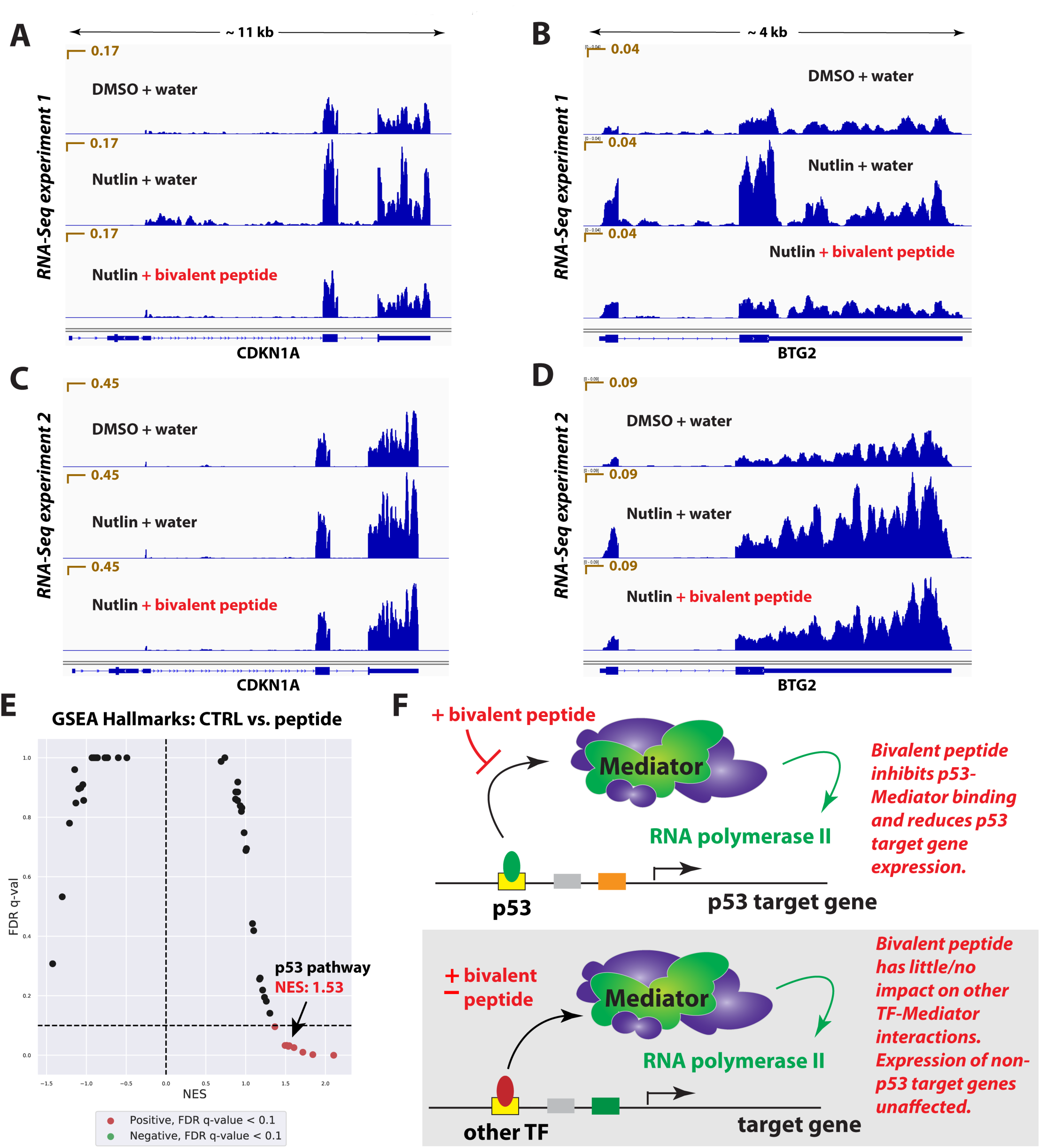
Bivalent peptide blocks p53 pathway activation but has minimal effect on non-p53 target genes. (*A, B*) Representative IGV traces at select p53 target gene loci, showing induction with Nutlin-3a and reduced induction in presence of bivalent peptide (RNA-Seq experiment 1). (*C, D*) Representative IGV traces from RNA-Seq experiment 2, showing similar but not identical effects compared with RNA-Seq experiment 1. (*E*) GSEA hallmarks moustache plot comparing unstimulated cells (i.e. DMSO) in presence or absence of bivalent peptide (RNA-Seq experiment 1 and experiment 2 combined). The plot reveals limited gene expression changes compared with Nutlin-3a treated cells (e.g. Fig. 3C). Heat map details on the modest p53 activation is shown in Supplemental Figure 6C, D. (*F*) Model; Mediator serves as a bridge between DNA-binding TFs and the pol II enzyme. The bivalent peptide effectively competes with p53 to prevent its binding to Mediator and subsequent activation of target genes *in vitro* and in cells. At non-p53 target genes, which are activated in part through other TF-Mediator interactions (via different Mediator subunits), the bivalent peptide has minimal impact on pol II transcription.

As expected, Nutlin-3a induced expression of p53 target genes (Fig. 3C), consistent with previous studies in HCT116 cells (Allen et al. 2014). Strikingly, however, Nutlin-induced activation of p53 target genes was diminished in cells treated with the bivalent peptide (vs. controls; Fig. 3D). Inhibition by the bivalent peptide was observed across a core set of p53 target genes (Andrysik et al. 2017), identified as p53-inducible across cell types (Fig. 3E).

A confounding issue with these experiments was that cellular uptake of the bivalent peptide could not be accurately measured and could potentially vary from experiment-to-experiment. To more thoroughly assess this possibility, we completed an additional round of RNA-Seq experiments in control vs. Nutlin-treated HCT116 cells (Supplemental Fig. 4B). These additional experiments (RNA-Seq experiment 2) were completed with a slightly modified protocol that included treatment in serum-free media as an effort to enhance peptide uptake (Brown et al. 2013; Wachter et al. 2017). Although the overall Nutlin response was reduced (GSEA p53 pathway NES = 2.76 in RNA-Seq experiment 1 vs. NES = 1.59 in RNA-Seq experiment 2), the data from RNA-Seq experiment 2 (biological triplicate; Supplemental Fig. 5) were consistent with the first series of biological replicates (RNA-Seq experiment 1; Fig. 3C**-E**). Specifically, Nutlin-induced activation of p53 target genes was inhibited in the presence of the bivalent peptide, providing further evidence that the bivalent peptide blocks p53 response in cells. However, the scope and magnitude of the effect was reduced in this second, independent set of RNA-Seq experiments (e.g. compare Fig. 3C, D & Supplemental Fig. 5C, D). This broadly consistent but variable magnitude effect can also be seen from RNA-Seq reads at specific p53 target genes (RNA-Seq experiment 1, Fig. 4A, B; RNA-Seq experiment 2, Fig. 4C, D), which illustrate 1) Nutlin induction and 2) inhibition of p53 target gene activation in the presence of the bivalent peptide. We interpret the variable magnitude of p53 suppression (RNA-Seq experiment 1 vs. RNA-Seq experiment 2) as a reflection of variable cellular uptake of the bivalent peptide.

### Bivalent peptide has modest transcriptional effect in absence of p53 activation

An expectation of our experimental strategy was that the bivalent peptide, which was designed based upon p53AD structure, would selectively block p53 function. Support for selective p53 inhibition was observed *in vitro*, upon comparison of the effects of the bivalent peptide on GAL4-p53AD vs. GAL4-VP16AD (Fig. 2C, D). RNA-Seq experiments in HCT116 cells represented a more rigorous test for selectivity. HCT116 cells express hundreds of sequence-specific, DNA-binding TFs, including high-level expression of TFs that define the cell lineage (Hnisz et al. 2013). For HCT116 cells, these TFs include SREBF1, ELF3, JUNB, NR2F1, and MYC. To assess the general impact of the bivalent peptide on pol II transcription, we compared RNA-Seq data from DMSO control cells (i.e. no Nutlin treatment) in the presence/absence of bivalent peptide. The data revealed that, in contrast to Nutlin-treated cells, the bivalent peptide had a minor effect on pol II transcription, genome-wide, in both sets of RNA-Seq experiments. For example, PCA plots showed clustering of DMSO control and peptide-treated samples compared with Nutlin-treated samples (Supplemental Fig. 6A, B), and heat maps derived from the RNA-Seq data showed modest changes in control vs. peptide-treated cells (Supplemental Fig. 6C, D). GSEA showed a reduced impact of the bivalent peptide on global transcription in the absence of Nutlin stimulation; however, enrichment of some hallmark gene sets was detected (Fig. 4E). Interestingly, evidence for p53 activation was observed in bivalent peptide-treated cells (vs. untreated controls) in RNA-Seq experiments 1 and 2, and modest p53 activation could be seen in heatmaps of p53 target genes (Supplemental Fig. 6C, D). This observation could reflect an ability of the bivalent peptide to mimic p53AD-Mediator binding to activate transcription (as suggested *in vitro* for the BP1.4 peptide, Supplemental Fig. 1C) and/or an ability to inhibit MDM2 and/or MDM4 binding to p53 in HCT116 cells. Overall, the RNA-Seq data from uninduced cells (i.e. not treated with Nutlin) were consistent with our *in vitro* results (Fig. 2) and suggest that the bivalent peptide does not broadly impact pol II transcription but can selectively block activation of p53 target genes in HCT116 cells (Fig. 4F).

A final comparison that could be made from the RNA-Seq data was to assess the effect of Nutlin-3a in peptide-treated cells. Given that the bivalent peptide alone could trigger a mild p53 response (Supplemental Fig. 6C, D), we asked whether Nutlin would still activate p53 target genes in this context. As expected, GSEA comparisons revealed that 1) Nutlin was able to induce p53 target gene expression even in the presence of the bivalent peptide, but 2) Nutlin induction of p53 target genes was reduced in this context (Supplemental Fig. 6E, F). For instance, the Normalized Enrichment Score (NES) for Nutlin-induced p53 pathway activation in the absence of bivalent peptide was 2.76 (experiment 1) or 1.59 (experiment 2); by contrast, the corresponding NES from peptide-treated cells (+ Nutlin) was 1.97 or 0.92, respectively. These results further support a role for the bivalent peptide in the suppression of p53 response in Nutlin-treated cells.

Although few TF-Mediator interactions have been characterized in detail, an emerging theme is the importance of bivalent or multi-valent interactions (Herbig et al. 2010; Currie et al. 2017). Our results demonstrate the importance of both p53 activation domains in Mediator-dependent transcription activation. A bivalent interaction may be required to selectively bind Mediator, as other proteins are bound by p53AD1 or p53AD2 individually (Lin et al. 1994; Ferreon et al. 2009) or can have their affinity enhanced by p53AD phosphorylation (Krois et al. 2016). The stapled, bivalent peptide is likely a poor substrate for site-specific phosphorylation, which may contribute to its selectivity and effectiveness in cells. However, we cannot exclude the possibility that other functionally relevant interactions are blocked by the bivalent peptide, which may contribute to its cellular activity.

Stapled peptides have shown promise as molecular therapeutics (Morrison 2018), including several that target p53 interactions with the MDM2 and/or MDM4 inhibitor proteins (Bernal et al. 2010; Brown et al. 2013; Chang et al. 2013; Carvajal et al. 2018). Our staple design for p53AD1 was based upon previous studies (Bernal et al. 2007) and enforces an α-helical conformation. The staple itself may also contribute to Mediator binding, and future experiments are needed to structurally define the interface targeted by the bivalent peptide. Although p53 normally functions as a tumor suppressor, gain-of-function p53 mutations are common and can be oncogenic (Freed-Pastor and Prives 2012; Kastenhuber and Lowe 2017); thus, blocking the activity of such p53 mutants is a viable therapeutic strategy (Brosh and Rotter 2009).

Historically, TFs have been intractable as therapeutic targets. Our proof-of-concept study suggests that desired transcriptional outcomes can be achieved by circumventing the TF entirely and targeting the human Mediator complex instead. Additional support for this concept was provided by Arthanari et al. (Nishikawa et al. 2016), in which the Med15-Pdr1 interaction was blocked with a small molecule in yeast (*C. glabrata*). Analogous to our results with p53, they showed that disruption of the Mediator-Pdr1 interaction prevented activation of Pdr1 target genes in yeast cells. Whereas the number of distinct, functionally relevant Mediator-TF interfaces is limited in yeast, TFs target many different sites on human Mediator (Poss et al. 2013). An implication is that blocking a single Mediator-TF interaction will not affect other signal-responsive or lineage-specific TFs (Fig. 4F), thus providing a means to selectively alter gene expression patterns. Our results suggest this strategy could be applied toward p53 and other human TF activation domains that target distinct Mediator subunits.

Funding support provided by the NCI (CA170741 to AS & DJT), Golfers Against Cancer (DJT), the NIA (F30AG054091 to CBL), and the NIH (GM117370 to DJT; GM125871 to RDD; T32GM008759 to BLA & JDR; F32GM122361 to TR).

Author contributions: BLA: biochemical and cellular assays; KQ: peptide design and synthesis; CBL: cell-based assays, RNA-Seq; JDR: RNA-Seq data analysis; TR: cell-based assays, RNA-Seq; RDD: RNA-Seq; Schepartz: designed and supervised research; DJT: designed and supervised research, wrote manuscript. All authors contributed to manuscript preparation.

## Data availability

RNA-Seq data have been uploaded on GEO: GSE135870.

## Materials and methods (*See also Supplemental Information*)

Purification of PIC factors for in vitro transcription

TFIIA, TFIIB, TFIID, TFIIE, TFIIF, TFIIH, Mediator, and pol II were purified as described (Knuesel et al. 2009).

### In vitro transcription

Chromatinized templates and *in vitro* transcription assays were generated and completed as described (Knuesel et al. 2009).

### HCT116 cell culture

HCT116 cells were grown in McCoy’s media (Gibco, 16600082) with Gibco 100x Antibiotic-Antimycotic (Fisher Sci, 15240062) penicillin-streptomycin and 10% fetal bovine serum (FBS) supplementation.

## Supplemental Information

**Supplemental Figure 1.**
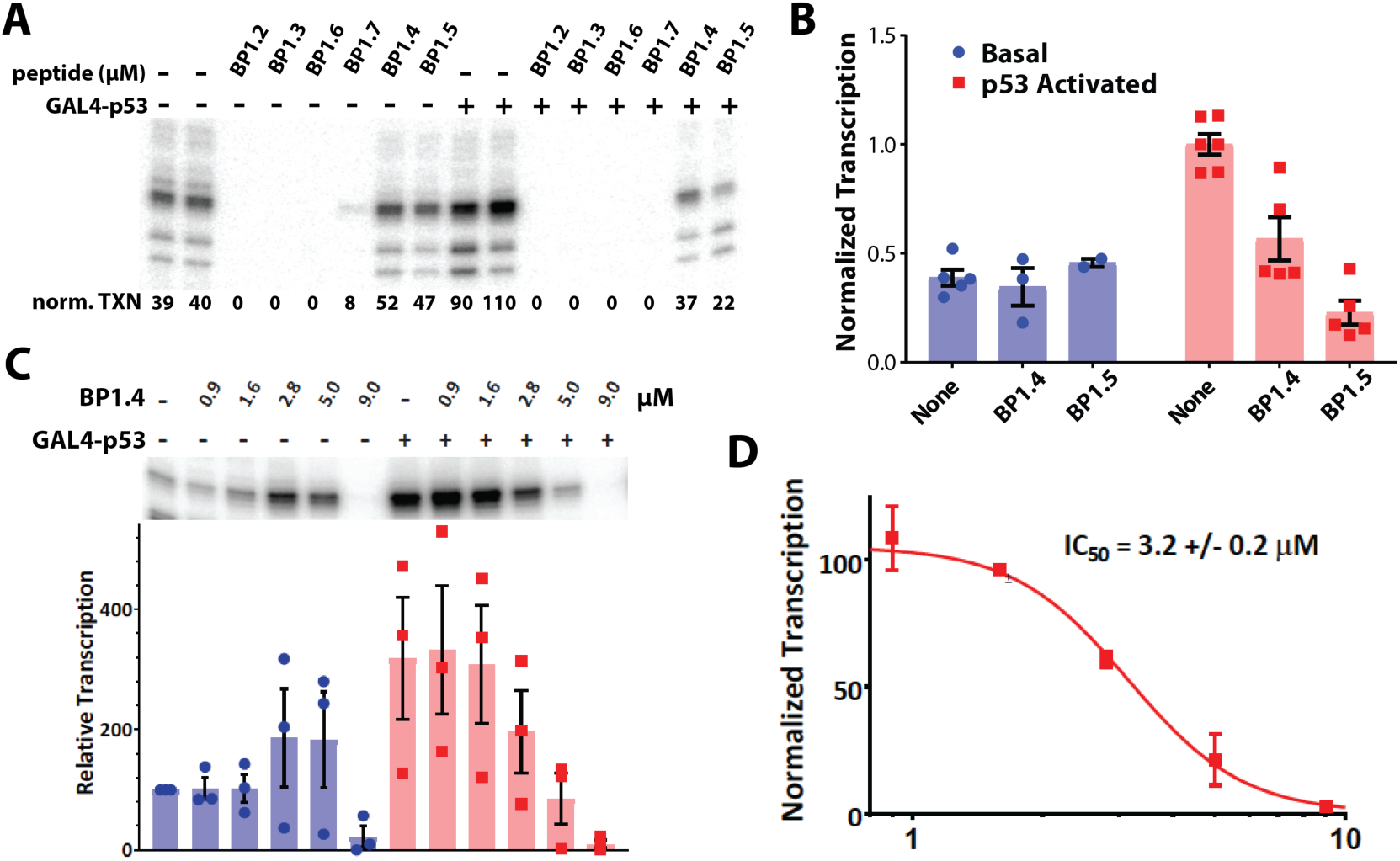
Functional screening of stapled and unstapled p53AD1 mimics. (*A*) Representative data (*in vitro* transcription) using each peptide at 5 *µ*M concentration, in presence (+) or absence (–) of GAL4-p53. Note that only BP1.4 and BP1.5 show ability to inhibit p53-activated transcription while not markedly affecting basal transcription. (*B*) Scatter plot summarizing *in vitro* transcription data for BP1.4 and BP1.5 peptides in absence (basal) or presence (activated) of GAL4-p53. (*C*) Representative data (top) and scatter plot (bottom) summarizing results from titration experiments with BP1.4 peptide under basal (– GAL4-p53) or activated (+ GAL4-p53) conditions. (*D*) IC_50_ plot summarizing inhibitory activity of BP1.4 peptide. For all data panels (*A-D*), transcription was normalized to GAL4-p53 in absence of added peptide.

**Supplemental Figure 2.**
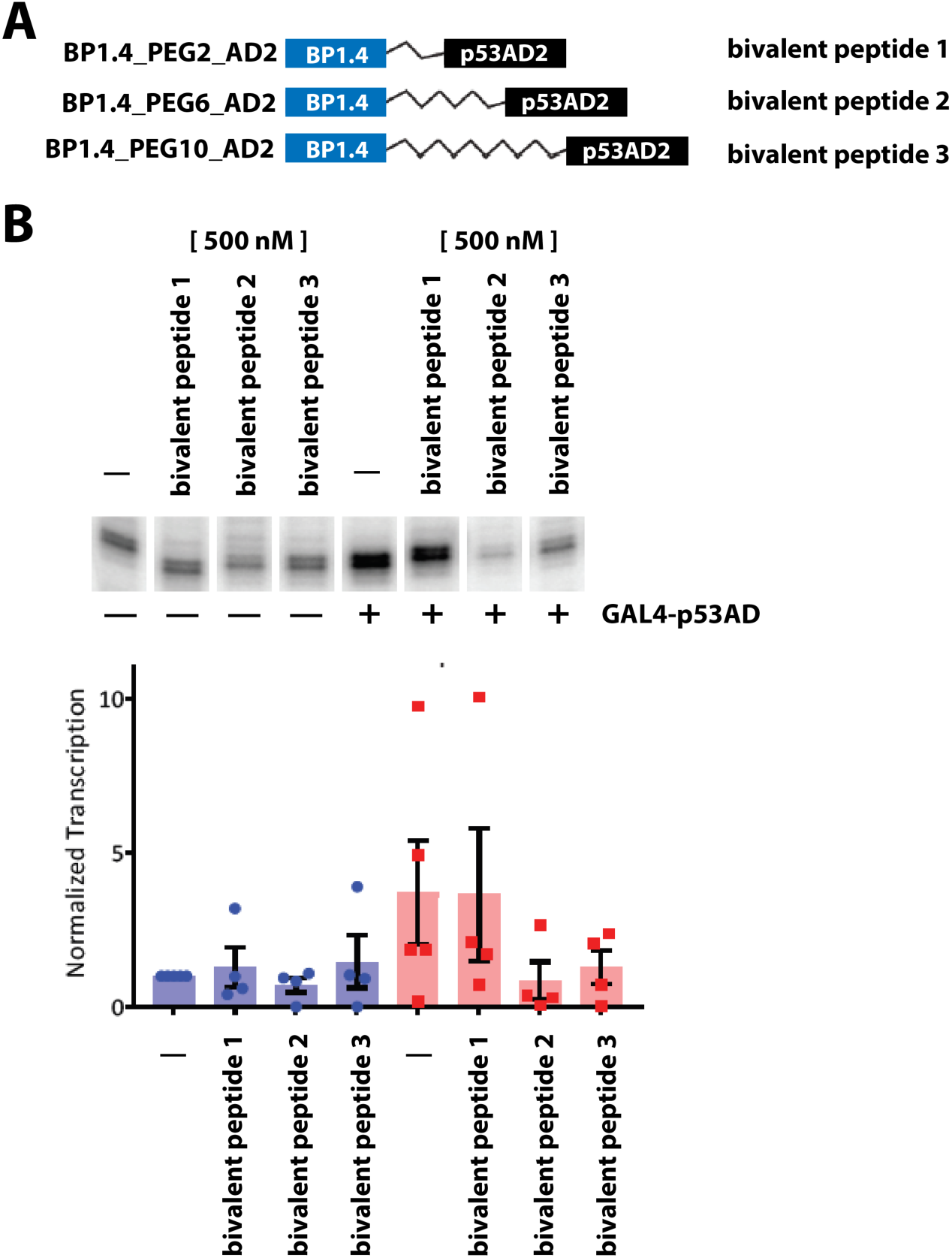
Testing different PEG linker lengths to tether BP1.4 (stapled p53AD1 mimic) to p53AD2 sequence. (*A*) Schematic of 3 different PEG linkers to generate bivalent peptide 1, 2, or 3. (*B*) Representative *in vitro* transcription data (top) and scatter plot summary of bivalent peptides with different PEG linker lengths. Note that a PEG6 or PEG10 linker showed enhanced ability to block p53-activated transcription, whereas PEG2 linker (i.e. bivalent peptide 1) did not.

**Supplemental Figure 3.**
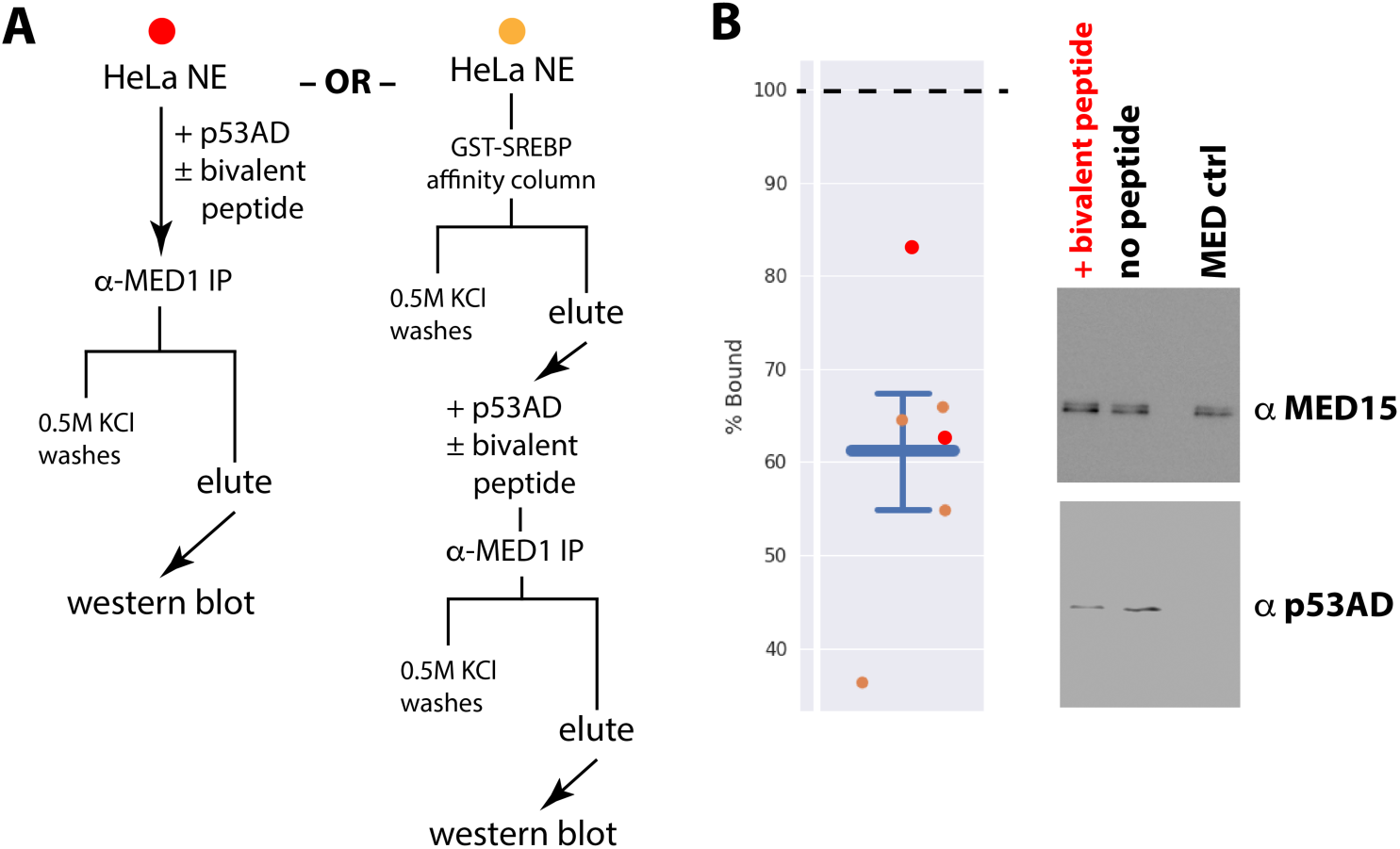
Bivalent peptide blocks p53AD-Mediator binding. (*A*) Overview of binding assays used. A crude Mediator sample was isolated from HeLa NE with a GST-SREBP affinity column (Naar et al. 1999) prior to incubation with p53AD ± bivalent peptide in one of the protocols (right). (*B*) Scatterplot (left) summarizing p53AD binding to Mediator in presence of bivalent peptide. The percent binding is relative to p53AD bound to Mediator in absence of added peptide; p53AD quantitation was normalized to total Mediator, as assessed by quantitation of MED15 signal (and/or MED1 in some cases; n = 4 biological and 6 total replicates). Red or orange dots correspond to data from binding protocol shown in panel *A*. Representative data (western blot) shown at right.

**Supplemental Figure 4.**
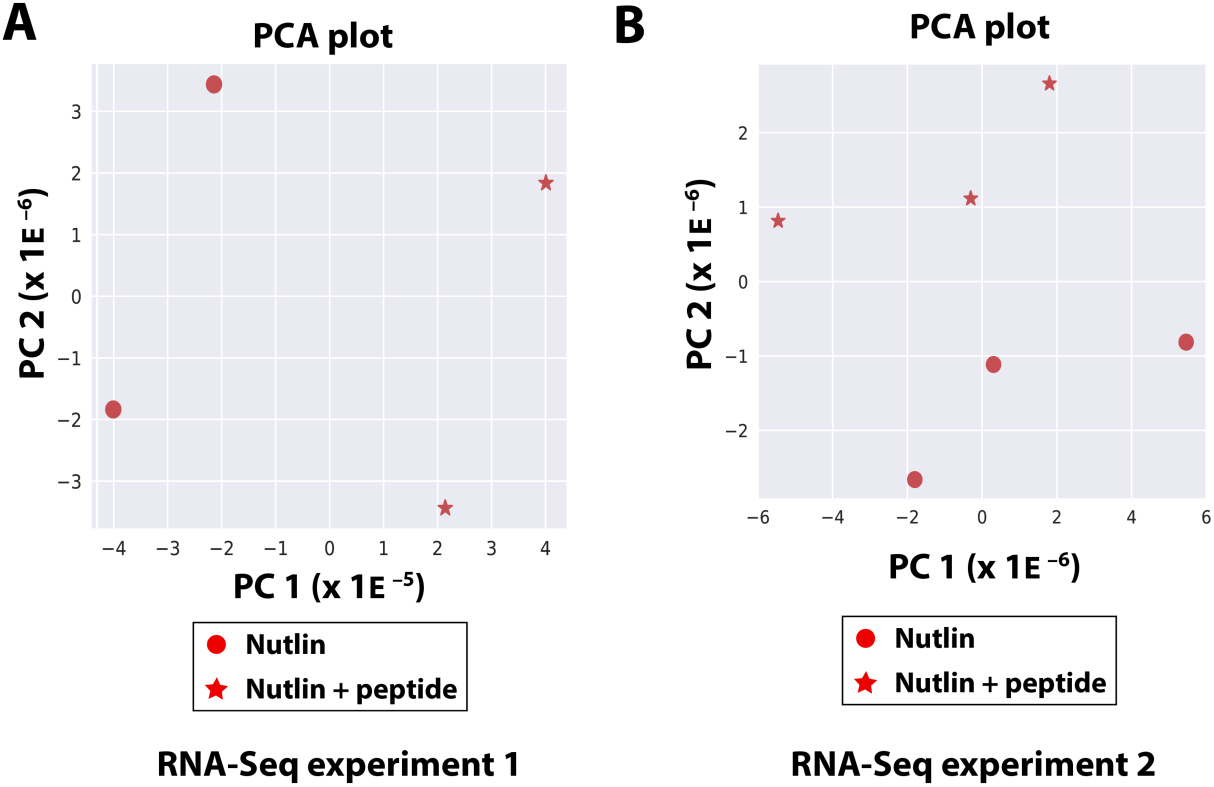
Principal Component Analysis for RNA-Seq experiment 1 & 2. (*A*) PCA plot for biological replicate RNA-Seq experiment 1. (*B*) PCA plot for biological triplicate RNA-Seq experiment 2.

**Supplemental Figure 5.**
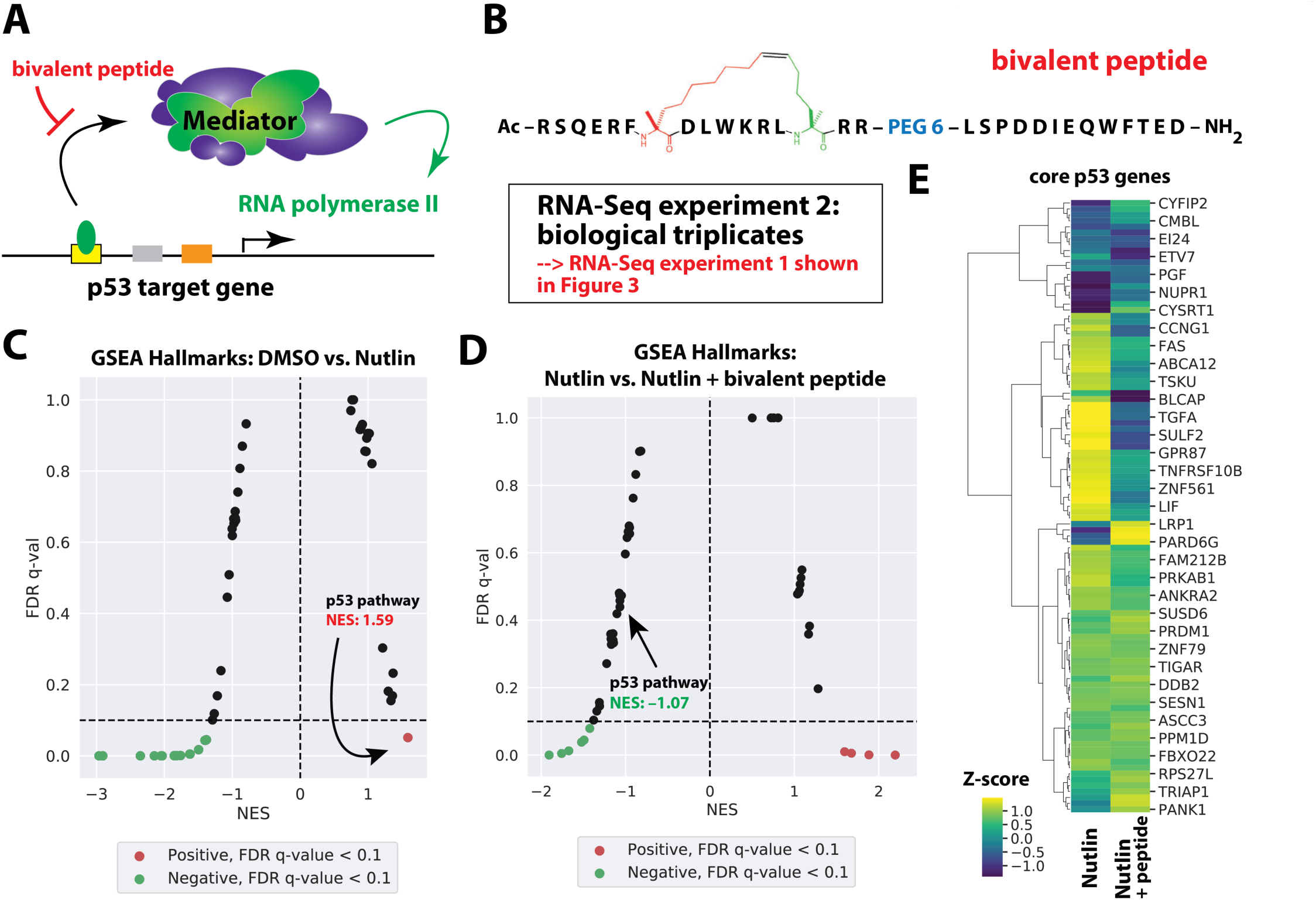
Bivalent peptide blocks p53 response in Nutlin-treated HCT116 cells (RNA-Seq experiment 2). (*A*) Simplified schematic showing Mediator recruitment and activation of pol II function via p53. (*B*) Structure of bivalent peptide. (*C*) GSEA hallmarks moustache plot showing p53 pathway activation upon treatment with Nutlin-3a (10 *µ*M, 3h). (*D*) GSEA hallmarks moustache plot showing effect of bivalent peptide in Nutlin-treated cells. Note the p53 pathway shows negative enrichment in cells treated with the bivalent peptide. (*E*) Heat map showing relative expression of core set of 103 p53 target genes (Andrysik et al. 2017) in control (DMSO) vs. Nutlin-treated cells, in absence or presence of bivalent peptide. In agreement with GSEA plots from panels C & D, Nutlin-3a induction of p53 target genes is reduced in cells treated with bivalent peptide.

**Supplemental Figure 6.**
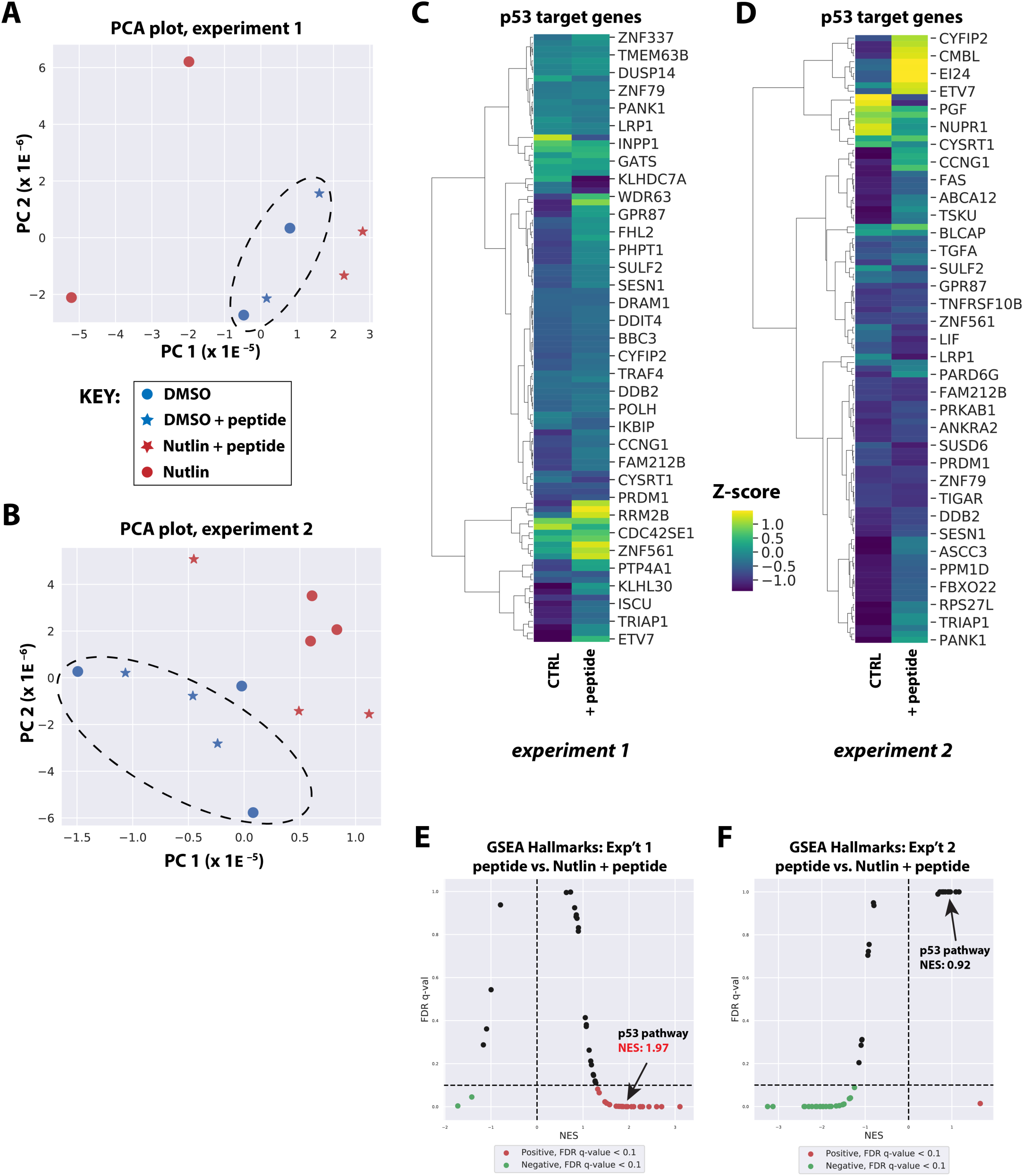
Bivalent peptide has minimal impact on HCT116 transcriptome in absence of p53 stimulation. Principal Component Analysis (PCA) for RNA-Seq experiment 1 (*A*) and RNA-Seq experiment 2 (*B*) for unstimulated HCT116 cells (i.e. DMSO) in the presence or absence of bivalent peptide. Clustering of DMSO and DMSO + peptide samples highlighted by enclosed circle. (*C, D*) Heat maps showing relative expression of core p53 target genes (Andrysik et al. 2017) in unstimulated (DMSO) HCT116 cells, in absence or presence of bivalent peptide. RNA-Seq experiment 1 (*C*) or RNA-Seq experiment 2 (*D*). (*E, F*) GSEA hallmarks moustache plots showing effect of Nutlin in the presence of bivalent peptide (RNA-Seq experiment 1 or 2, respectively). Note the p53 pathway is still induced by Nutlin in the presence of bivalent peptide, but the magnitude of the induction is reduced compared with Nutlin + no peptide experiments (e.g. GSEA plot in Fig. 3C or Supplemental Fig. 5C).

**Supplemental Table 1.** Summary of GSEA results for RNA-Seq experiments 1 & 2.

## Supplemental Materials and methods

### Radiolabeling of the reverse transcription primer

A reverse transcriptase (RT) primer was synthesized to complement the RNA transcript 85 bases downstream of the transcription start site. The RT primer was radiolabeled in polynucleotide kinase (PNK) buffer (70 mM Tris-HCl pH 7.6, 10 mM MgCl_2_, 5 mM DTT) with the addition of about 150 µCi [γ-^32^P]ATP, 6 U of T4 PNK and 48 ng of the RT primer in a final volume of 10 *µ*L. The reactions were then incubated at 37 °C for 45 minutes. A glycogen mixture (10 mM Tris pH 7.5, 34 mM EDTA, 1.33 mg/mL glycogen) was then added to bring the volume to 25 *µ*L, and the reaction was passed through a G-25 column to remove excess free [γ-^32^P]ATP. An additional 25 *µ*L of TE buffer (10 mM Tris pH 7.5, 1 mM EDTA) was added. The radiolabeled primer was then stored at 4 °C until needed (up to 1 week).

### In vitro transcription

Chromatinized templates and *in vitro* transcription assays were generated and completed as described (Knuesel et al. 2009). Briefly, each activator (GAL4-p53AD or GAL4-VP16AD) was titrated to yield maximum transcription. While the activator bound the template, the general transcription factors (GTFs) were mixed in 0.1 M HEMG (10 mM HEPES pH 7.6, 100 mM KCl, 0.1 mM EDTA, 10% glycerol, 5.5 mM MgCl_2_) to give approximate final concentrations of 40 nM TFIIA, 10 nM IIB, 0.8 nM TFIID, 10 nM TFIIE, 10 nM TFIIF, 0.5 nM TFIIH and 2 nM pol II. A non-limiting amount of Mediator was then diluted in a separate salts mix (10 mM HEPES pH 7.6, 100 mM KCl, 2.5% PVA, 2.5% PEG, 7.5 mM MgCl_2_), along with 400 U of RNAseOUT, about 300 ng PC4 and about 300 ng HMGB1. On ice, the desired concentration of peptide was then added to the Mediator mix, followed by the GTF mix at a 5:11 ratio. The GTFs, Mediator and peptide were then incubated at least 5 minutes at 30 °C. Then, 15 *µ*L of the mixture was added to each reaction. PIC assembly proceeded for 15 minutes, then transcription was initiated by adding 5 *µ*L of a solution containing 5 mM of each NTP. After thirty minutes, reactions were stopped with the addition of 150 *µ*L Stop Buffer (20 mM EDTA, 200 mM NaCl, 1% SDS, 100 µg/mL Proteinase K, 100 µg/mL glycogen) and incubating at 37 °C for 15 minutes. RNA was isolated with 100 *µ*L phenol/chloroform/isoamyl alcohol (pH 7.7-8.3); 140 *µ*L of the aqueous phase was mixed with 5 *µ*L, 7.5 M ammonium acetate and 5 *µ*L of twenty-fold diluted, radiolabeled (^32^P) Reverse Transcriptase (RT) probe and transferred to a 500 *µ*L microfuge tube. The RNA was then precipitated by adding 375 *µ*L, 100% cold ethanol and placing at −20 °C for at least an hour.

### Primer extension

Reactions were spun down at 14K RPM for 20 minutes and the ethanol was removed. Pellets were then briefly dried (speedvac) and resuspended in 10 *µ*L Annealing Buffer (10 mM Tris-HCl pH 7.8, 1 mM EDTA, 250 mM KCl). The resuspended RNA was then incubated in a thermocycler as follows: 85 °C for 2 minutes, cool to 58 °C at 30 sec/degree, 58 °C for 10 minutes, 57 °C for 20 minutes, 56 °C for 20 minutes, 55 °C for 10 minutes, and cool to 25 °C at 30 seconds/degree. 38 *µ*L of RT mix (20 mM Tris-HCl pH 8.7, 10 mM MgCl_2_, 0.1 mg/mL actinomycin D, 330 *µ*M of each dNTP, 5 mM DTT, 0.33 U/*µ*L Moloney Murine Leukemia Virus (MMLV) reverse transcriptase) was then added to the annealing reactions and allowed to extend for forty-five minutes at 37 °C. Reactions were then stopped and precipitated by adding 300 *µ*L cold ethanol and placed at −20 °C for at least an hour.

### Separation and visualization of transcript cDNA by denaturing polyacrylamide gel electrophoresis

The cDNA reactions were spun down at 14K RPM for 25 minutes and the ethanol was removed from the pellets. After briefly drying pellets (speedvac), cDNA was resuspended in 6 *µ*L formamide loading buffer (75% formamide, 4 mM EDTA, 0.1 mg/mL xylene cyanol, 0.1 mg/mL bromophenol blue, 33 mM NaOH), heated for 3 minutes at 90 °C and loaded onto a denaturing polyacrylamide gel (89 mM Tris base, 89 mM boric acid, 2 mM EDTA, 7 M Urea, 6% acrylamide/bisacrylamide [19:1]). Gels were run at 35 W for about 1.5 hours, then removed on filter paper and dried for 1 hour at 80 °C. Gels were then exposed on a phosphorimager screen.

### Peptide Synthesis Reagents

All purchased reagents were used without further purification. Standard Fmoc-protected amino acids were purchased from Novabiochem (San Diego, CA). Fmoc-protected olefinic amino acids, (*S*)-N-Fmoc-2-(4’-pentenyl)alanine and (R)-N-Fmoc-2-(7’-octenyl)alanine, were purchased from Okeanos Tech Jiangsu Co., Ltd (Jiangsu, P.R. China). Rink amide resin, *N,N*-dimethylformamide (DMF), *N*-hydroxybenzotriazole (HOBt), and Grubbs Catalyst™ 1^st^ Generation were purchased from Sigma-Aldrich (St. Louis, MO). Trifluoroacetic acid (TFA) and dichloroethane (DCE) were purchased from Acros Organics (Fair Lawn, NJ). *N,N,N’,N’*-tetramethyl-uronium-hexafluoro-phosphate (HBTU) and diisopropylethylamine (DIEA) were purchased from AmericanBio (Natick, MA). Anhydrous piperazine and 6-chlorobenzotriazole-1-yloxy-tris-pyrrolidinophosphonium hexafluorophosphate (PyClocK) was purchased from EMD Millipore (Billerica, MA). Acetic anhydride was purchased from ThermoScientific, Pierce Biotechnology (Rockford, IL).

### Solid Phase Peptide Synthesis

Peptides were synthesized using standard Fmoc chemistry with Rink amide resin on Biotage® Initiatior+ Alstra from Biotage (Charlotte, NC) using microwave acceleration. Fmoc deprotections were performed using 5% piperazine with 0.1 M HOBt to reduce aspartimide formation in DMF. Coupling reactions were performed using 5 equivalents of amino acid, 4.9 equivalents of HBTU, 5 equivalents of HOBt, and 10 equivalents of DIEA in DMF at 75°C for 5 min. Fmoc-NH-(PEGn)-COOH linkers were coupled as amino acids were. All arginine residues were double coupled at 50°C. Olefinic 55 side-chain bearing residues were coupled using 3 equivalents of amino acid, 3 equivalents of PyClocK, and 6 equivalents of DIEA and stapled for 2 hours at room temperature PyClocK. Residues following olefinic residues were double coupled using standard coupling procedures. N-terminally capped peptides were generated by treating Fmoc-deprotected resin with 100 equivalents acetic anhydride and 100 equivalents DIEA for 10 minutes at room temperature. Following synthesis, resin was washed thoroughly with alternating DMF (5 mL) and DCM (10 mL) washes before subsequent cyclizing, labeling, and cleavage.

### Ring Closing Olefin Metathesis

Peptides containing olefinic amino acids were washed with DCM (3 x 1 min) and DCE (3 x 1 min) prior to cyclizing on resin using Grubbs Catalyst I (20 mol % compared to peptide, or 1 equivalent compared to resin) in DCE (4 mL) for 2 h under N2. The cyclization step was performed twice (Kim et al. 2011). The resin was then washed three times with DCM (5 mL) before washing with MeOH (5 mL x 5 min) twice to shrink the resin. The resin was dried under a stream of nitrogen overnight.

### Peptide Cleavage

After shrinking and drying overnight, the peptide was cleaved from the resin using a 3 mL solution of trifluoroacetic acid (TFA) (81.5%), thioanisole (5%), phenol (5%), water (5%), ethanedithiol (EDT) (2.5%) and triisopropylsilane (TIPS) (1%) for 2 hours at RT on an orbital shaker. Cleaved peptides were precipitated in diethyl ether (40 mL, chilled to −80°C), pelleted by centrifugation, washed with additional diethyl ether (40 mL, −80°C), pelleted, redissolved in a solution of acetonitrile (ACN) and water (15% CAN), frozen, lyophilized to dryness, and reconstituted in 1 mL dimethyl sulfoxide (DMSO) prior to purification by high-performance liquid chromatography (HPLC).

### Peptide Purification by HPLC

Peptide solutions were filtered through nylon syringe filters (0.45 μm pore size, 4 mm diameter, Thermo Fisher Scientific) prior to HPLC purification. Peptides were purified using an Agilent 1260 Infinity HPLC system on a reverse phase Triaryl-C18 column (YMC-Triaryl-C18, 150 mm x 10 mm, 5 μm, 12 nm) (YMC America, Inc.) over H_2_O/ACN gradients containing 0.1% TFA. Peptides were detected at 214 nm and 280 nm. Peptide purity was verified using a Shimadzu Analytical ultra-performance liquid chromatography (UPLC) system (ES Industries, West Berlin; Shimadzu Corporation, Kyoto, Japan) and a C8 reverse phase (Sonoma C8(2), 3 μm, 100 Å, 2.1 x 100 mm) analytical column. Analytical samples were eluted over a gradient of 15-57 60% ACN in water containing 0.1% TFA over 15 min with detection at 214 and 280 nm.

### In vitro binding assays

Starting from 180 *µ*L HeLa nuclear extract (which contains Mediator), bivalent peptide was added (to 5*µ*M concentration) followed by addition of purified p53AD (residues 1-70; to 2 *µ*M concentration). A parallel experiment lacked added bivalent peptide. Each sample was allowed to incubate, with mixing, for 2 hours at 4°C. Each sample was then incubated, with mixing, over an anti-MED1 affinity resin (to immunoprecipitate Mediator from the sample) for 90 minutes at 4°C. The resin was then washed 4 times with 20 resin volumes with 0.5M KCl HEGN (20 mM HEPES, pH 7.9; 0.1 mM EDTA, 10% glycerol, 0.1% NP-40) and once with 0.15M KCl HEGN (0.02% NP-40). Material that remained bound to the resin (i.e. Mediator) was eluted with 1M glycine, pH 2.2 and subsequently probed by western. As an alternate protocol, HeLa nuclear extract (1 mL) was first incubated over a GST-SREBP affinity column, washed 5 times with 0.5M HEGN, once with 0.15M HEGN, and eluted with 30 mM glutathione buffer, as described (Ebmeier and Taatjes 2010). This material (160 *µ*L), which is enriched in Mediator, was then incubated with p53AD (residues 1-70; to 2 *µ*M concentration) in the presence or absence of bivalent peptide (5 *µ*M) at 4°C for 1 hour. Then each sample was incubated, with mixing, over an anti-MED1 affinity resin, washed, eluted, and probed by western as described above.

### Experimental time frame for RNA-Seq experiments

In a series of experiments in SJSA cells, we initially tested whether the bivalent peptide could cause a phenotypic change. SJSA cells are unusually sensitive to Nutlin-3a (Vassilev et al. 2004) and therefore if the p53 response could be persistently blocked by the bivalent peptide, peptide-treated cells would show enhanced survival following Nutlin treatment. Starting with a 24-hour Nutlin treatment (10 *µ*M), we observed no significant effect of the bivalent peptide: similar percentages of cell death were observed in control vs. peptide-treated populations as analyzed by CellTiter-Glo assay (Promega). Although these results could be attributed to poor cellular uptake of the bivalent peptide, we also suspected that the peptide was active in cells for only a limited time (e.g. before being secreted or degraded). We next determined that a 6-hour Nutlin treatment time was the shortest that would still trigger significant SJSA cell death within 24-48 hours. However, we obtained similar results with 6-hour (or 12-hour) Nutlin treatment times (± bivalent peptide). We then tested the prospect of RNA-Seq experiments, in hopes that gene expression changes and shorter time frames would allow an assessment of bivalent peptide effects. Here, we used HCT116 cells, which show strong transcriptional response to Nutlin (Allen et al. 2014). For RNA-Seq experiments, we needed a time frame long enough to allow accumulation of p53 target gene mRNAs but short enough to enable maximum activity of the bivalent peptide (e.g. prior to its secretion, export, and/or degradation). Using RT-qPCR assays, we confirmed that a 3-hour Nutlin treatment was a minimum amount of time to reliably detect induction of p53 target genes. In addition, parallel assays confirmed that the bivalent peptide was blocking activation of p53-target genes in HCT116 cells during this time frame.

### Electroporation of bivalent peptide into HCT116 cells (RNA-Seq experiment 1)

Two 6-well plates (HCT116 cells) were grown to about 80% confluency. Cells were then trypsinized, washed with PBS, and resuspended in 150 *µ*L Neon Buffer R. The cells were then split into two groups: No peptide and 10 *µ*M peptide. The cells were then drawn into a 10 *µ*L Neon pipet tip, electroporated and ejected into 2 mL of McCoy’s 5A media without antibiotic. For each experiment, two cell electroporation aliquots were added to media containing either 0.1% DMSO (control) or 10 *µ*M Nutlin-3a (in DMSO, to a final concentration of 0.1%). For wells containing cells electroporated with peptide, an additional 200 nM peptide was added to the well to allow for peptide uptake during the experiment. The 6-well plate was then placed back at 37 °C for 3 hours. After 3 hours, cells were scraped from the plates, transferred to a 15 mL conical vial, pelleted at 1,000 x g, and washed in 10 mL phosphate buffered saline (PBS) solution. To isolate the nuclei, cells were resuspended in 10 mL lysis buffer (10 mM Tris-HCl, pH 7.4, 2 mM MgCl_2_, 3 mM CaCl_2_, 0.5% NP-40, 10% glycerol) and thoroughly mixed. The nuclei were spun down at 1,000xg for 10 minutes, the lysis buffer was removed, and 1 mL of TRIzol was added. The nuclear RNA was isolated as described in the TRIzol instructions, except an additional phenol/chloroform extraction and chloroform-only extraction were performed to reduce contaminants. RNA was precipitated and washed twice with 75% ethanol to further remove contaminants. The RNA was then converted to cDNA using the High Capacity cDNA kit from Thermo Fisher Scientific.

### Electroporation of bivalent peptide into HCT116 cells (RNA-Seq experiment 2)

One 15cm plate (HCT116 cells) was grown to about 70% confluency. Cells were then trypsinized, washed with PBS, and resuspended in 40 *µ*L Neon Buffer R. The cells were then split into two groups: No peptide and 10 *µ*M peptide. The cells were then drawn into a 10 *µ*L Neon pipet tip, electroporated and ejected into 2 mL of serum-free McCoy’s 5A media without antibiotic. For each experiment, two cell electroporation aliquots were added to media containing 0.1% DMSO (control) or 10 *µ*M Nutlin-3a (in DMSO, to a final concentration of 0.1%). For wells containing cells electroporated with peptide, an additional 200 nM peptide was added to the well to allow for peptide uptake during the experiment. The 6-well plate was then placed back at 37 °C for 3 hours. After 3 hours, cells were scraped from the plate, transferred to 2 mL eppendorf tubes, pelleted at 1,000 x g, and washed in 1 mL cold phosphate buffered saline (PBS) solution. To isolate the nuclei, cells were resuspended in 0.5 mL lysis buffer (10 mM Tris-HCl, pH 7.4, 2 mM MgCl_2_, 3 mM CaCl_2_, 0.5% NP-40, 10% glycerol) and mixed by pipetting up and down 20 times. The nuclei were spun down at 1,000 x g for 10 minutes, the lysis buffer was removed, and 200 *µ*L of TRIzol was added. The RNA was isolated as described in the TRIzol instructions, except an additional phenol/chloroform extraction and chloroform-only extraction were performed. RNA was precipitated and washed twice with 75% ethanol. The RNA was then converted to cDNA using the High Capacity cDNA kit from Thermo Fisher Scientific.

### RNA-Seq

Quality control was performed on raw reads using FastQC (Accessed 7/18/2019; https://www.bioinformatics.babraham.ac.uk/projects/fastqc/). Raw reads were then trimmed using bbduk (Accessed 7/18/2019; https://jgi.doe.gov/data-and-tools/bbtools/bb-tools-user-guide/bbduk-guide/) with options: ktrim=r qtrim=10 k=23 mink=11 hdist=1 maq=10 minlen=25 tpe tbo literal=AAAAAAAAAAAAAAAAAAAAAAA. Trimmed reads were then mapped to the human genome (hg38) using HISAT2 (Kim et al. 2015) (options: --very-sensitive). Mapped reads over genes were counted using bedtools multibamcov and RPKM normalized. Batch correction and PCA analysis was performed in R using limma (Ritchie et al. 2015). Heatmaps were generated in Python using seaborn (Accessed 7/18/2019; https://seaborn.pydata.org/introduction.html) with batch corrected counts from limma. Heatmap z-scores were calculated across treatments (rows) using all data relevant to each experiment with replicates either combined or separated where appropriate. Heatmap rows and columns were clustered using Ward’s method.

### Principal Component Analysis

PCA was performed using the standard prcomp function provided by the sva package for the R programming language (Leek et al. 2019). Batch effects from replicates completed on different days replicates were corrected using the removeBatchEffect function provided by the limma package (Ritchie et al. 2015) from the R programming language.

### Differential Expression analysis

Differential expression analysis was performed using the DESeq2 package (Love et al. 2014) for the R programming language. Counts were generated using the utility featurecounts. Initial analysis using counts across the full annotated gene showed significant skew, indicating that the baseline assumptions of the differential expression model did not hold. To correct, counts in the region from +500 of the TSS to −500 from the TES (Transcription End Site) were used to obtain suitable model weights. Those model weights were then used when performing differential expression across the full gene, which corrected the skew effect.

### Gene Set Enrichment Analysis

GSEA (Subramanian et al. 2005) was performed with the Broad Institute’s GSEA software on the GenePattern Server using the pre-ranked module. Normalized (DE-Seq2) (Love et al. 2014) log(2) fold-change values were used as the rank metric for all genes and compared against the Hallmark gene sets database for enrichment.

